# Seesaw Conformations of Npl4 in the Human p97 Complex and the Inhibitory Mechanism of a Disulfiram Derivative

**DOI:** 10.1101/2020.05.18.103085

**Authors:** Man Pan, Qingyun Zheng, Yuanyuan Yu, Huasong Ai, Yuan Xie, Xin Zeng, Chu Wang, Lei Liu, Minglei Zhao

**Affiliations:** Department of Biochemistry and Molecular Biology, The University of Chicago, Chicago, IL 60637, USA; Tsinghua-Peking Center for Life Sciences, Department of Chemistry, Tsinghua University, Beijing 100084, China; Peking-Tsinghua Center for Life Sciences, College of Chemistry and Molecular Engineering, Peking University, Beijing 100871, China

**Author notes:** Corresponding authors: Minglei Zhao, Lei Liu. These authors contributed equally to the work (Man Pan, Qingyun Zheng, Yuanyuan Yu).

## Abstract

p97, also known as valosin-containing protein (VCP) or Cdc48, plays a central role in cellular protein homeostasis^1^. Human p97 mutations are associated with several neurodegenerative diseases^2,3^. Targeting p97 and its cofactors is a strategy for cancer drug development^4^. Despite significant structural insights into the fungal homolog Cdc48^5–7^, little is known about how human p97 interacts with its cofactors. Recently, the anti-alcohol abuse drug disulfiram was found to target cancer through Npl4, a cofactor of p97^8^, but the molecular mechanism remains elusive. Here, using single-particle cryo-electron microscopy (cryo-EM), we uncovered three Npl4 conformational states in complex with human p97 before ATP hydrolysis. The motion of Npl4 results from its zinc finger motifs interacting with the N domain of p97, which is essential for the unfolding activity of p97. In vitro and cell-based assays showed that under oxidative conditions, the disulfiram derivative bis-(diethyldithiocarbamate)-copper (CuET) inhibits p97 function by releasing cupric ions, which disrupt the zinc finger motifs of Npl4, locking the essential conformational switch of the complex.

## INTRODUCTION

p97 (also known as valosin-containing protein (VCP) or Cdc48 in yeast) is a highly abundant cytoplasmic protein in eukaryotic cells^9^. It belongs to the protein family known as ATPase associated with diverse cellular activities (AAA+ protein superfamily)^10^. p97 has two tandem ATPase domains named D1 and D2 and an additional N domain at the N-terminus (**Fig. 1a**). Similar to many AAA+ proteins, p97 converts chemical energy from ATP hydrolysis to mechanical forces, which then relocate or unfold ubiquitinated substrates^11^. In vivo, p97 functions as a homomeric hexamer, a 540 kDa molecular machine with twelve copies of ATPase domains organized into two rings^12^. p97 plays a central role in cellular proteostasis. More than thirty mutations of human p97 have been discovered, and these are associated with a number of neurodegenerative diseases, including inclusion body myopathy, frontotemporal dementia, and familial amyotrophic lateral sclerosis^3^. Targeting p97 to disrupt cellular proteostasis is also a strategy for cancer therapy^4^.

**Figure 1:**
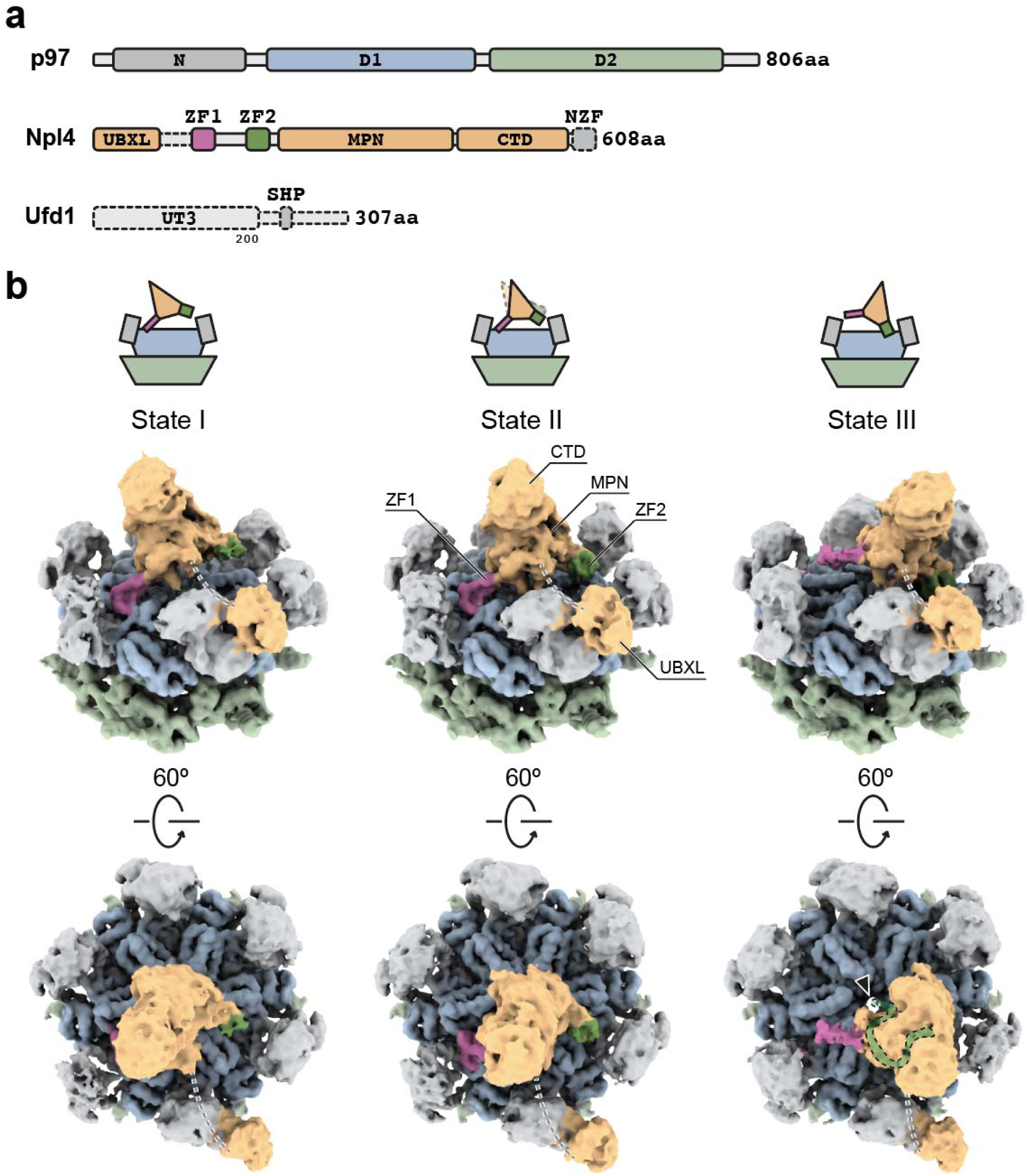
Three conformational states of human p97 in complex with Npl4/Ufd1. **a**, Domain architecture of human p97, Npl4 and Ufd1. Unresolved parts are shown with dotted lines. **b**, Single-particle cryo-EM maps (unsharpened) of human p97 in complex with Npl4/Ufd1. Three conformational states were resolved. The position of Npl4 relative to p97 is illustrated in the cartoons. Unresolved linkers between UBXL and ZF1 are represented by dotted lines. The central pore of the D1 ring in State III is highlighted by an arrowhead. The conserved groove of Npl4 that was found to bind an unfolded ubiquitin in the yeast complex structure^5^ was marked by a green band in State III.

One unique feature of p97 is that it can interact with a remarkable diversity of cofactors to participate in a variety of cellular pathways^13^. Most of these cofactors engage ubiquitinated substrates. The most versatile cofactor is a heterodimer consisting of two proteins: Npl4 and Ufd1 (denoted Npl4/Ufd1), which are likely involved in more than half of the cellular processes in which p97 participates^9^. A well-studied example is endoplasmic reticulum-associated degradation (ERAD), in which Cdc48, the yeast homolog of p97, recruits Npl4/Ufd1 to extract polyubiquitinated misfolded membrane proteins from the ER membrane, then processes and delivers the extracted proteins to the proteasome for degradation^14,15^. In particular, the FDA-approved drug disulfiram (tetraethylthiuram disulfide, trade name Antabuse), which has been used for a long time in the treatment of alcoholism, has recently been discovered to be effective against various cancer types in preclinical studies by targeting the cofactor Npl4^8^.

Recently, the cryo-electron microscopy (cryo-EM) structures of the thermophilic fungus Cdc48-Npl4/Ufd1 complex^6^ and the yeast Cdc48-Npl4/Ufd1-Eos complex^5^ were reported. In both structures, Npl4 interacts with one of the N domains of Cdc48 through its ubiquitin regulatory X-like (UBXL) domain and anchors the Mpr1/Pad1 N-terminal (MPN) domain on top of the D1 ring using both zinc finger motifs. One surprising finding was that the yeast Cdc48-Npl4/Ufd1 complex initiates substrate processing by unfolding one ubiquitin molecule in the absence of ATP binding or hydrolysis. The unfolded ubiquitin molecule binds to the groove of Npl4 and projects its N-terminal segment through both the D1 and D2 rings. The structure provided unprecedented insights into the substrate processing of Cdc48; however, it also raised a new question of how ubiquitin is unfolded and inserted into the central pore of the Cdc48 complex. More information about conformational dynamics is essential for a better understanding of this process.

Here, we investigated how human p97 interacts with its cofactor, Npl4/Ufd1, in the presence and absence of ubiquitinated substrates using cryo-EM and single-particle analyses. Our results showed that unlike the yeast homolog Cdc48, human p97 interacts with the cofactor Npl4/Ufd1 and its substrate in multiple conformational states without ATP hydrolysis. Furthermore, we revealed that the disulfiram derivative bis-(diethyldithiocarbamate)-copper (CuET) inhibits the unfolding activity of p97 and locks the conformational changes between p97 and Npl4/Ufd1 by releasing cupric ions under oxidative conditions.

## RESULTS

### Three conformational states of human p97 in complex with Npl4/Ufd1

Our work started with the structure determination of p97 in complex with the cofactor Npl4/Ufd1 to reveal the difference between the yeast and human systems. We determined the complex structure in the presence of ATPγS using cryo-EM and single-particle analyses (**Supplementary Fig. 1**). Using 3D classification, we resolved three different conformational states of the complex. In contrast to the complex structure of fungal Cdc48, in which Npl4 binds to the D1 ring using both zinc finger motifs (**Supplementary Fig. 2c**), human Npl4 binds to the D1 ring of p97 with only one zinc finger motif (**Fig. 1b and Supplementary Fig. 1**). In States I and II, Npl4 binds to the D1 ring of p97 using zinc finger motif 1 (ZF1, purple), whereas in State III, Npl4 binds to the same hydrophobic groove of the D1 ring using zinc finger motif 2 (ZF2, green). In both cases, the zinc finger motifs form hydrogen bonds with the backbone of an existing β-strand (F265-I269) in the large subunit of the D1 domain (**Supplementary Fig. 2a and b**). The difference between States I and II is that the main structure of Npl4, namely, the MPN and carboxyl-terminal domain (CTD), is rotated by ∼ 8 ° and is displaced by ∼ 13 Å with ZF1 as the fulcrum. Compared with States I and II, the main structure of Npl4 in State III is displayed by ∼ 49 Å and ∼ 38 Å, respectively (**Supplementary Fig. 2d**). Notably, when focused on the conserved Npl4 groove, we observed that only the center hole of p97 in State III is aligned with the groove and exposed in the top view (**Fig. 1b**); this was not observed in the complex structure of yeast Cdc48. Together, the conformational changes of these three states present a seesaw-like motion of Npl4 on top of the D1 ring. In all three states, the other zinc finger motif that is not interacting with the D1 ring is close to one of the N domains of p97, suggesting potential interactions that are observed in the complex structures with a polyubiquitinated substrate (**Fig. 2**). In addition to ZF1 and ZF2, a third zinc finger motif (NZF), which does not exist in the fungal homolog, is located at the very C-terminus of Npl4 (**Fig. 1a**). We were not able to resolve NZF due to the degradation of the local resolution (**Supplementary Fig. 1a**). A homology model of Npl4 was built based on the crystal structure of the thermophilic fungal homologue^6^. p97 adopts a six-fold symmetrical conformation in all three states. In fact, when applying a mask around the D1 and D2 rings of p97 and imposing a C6 point-group symmetry, the resolution of the 3D reconstruction improved to 2.8 Å, which allowed de novo model building (**Supplementary Figs. 1b and 2e-f**). Ufd1 was not resolved in the cryo-EM maps.

**Figure 2:**
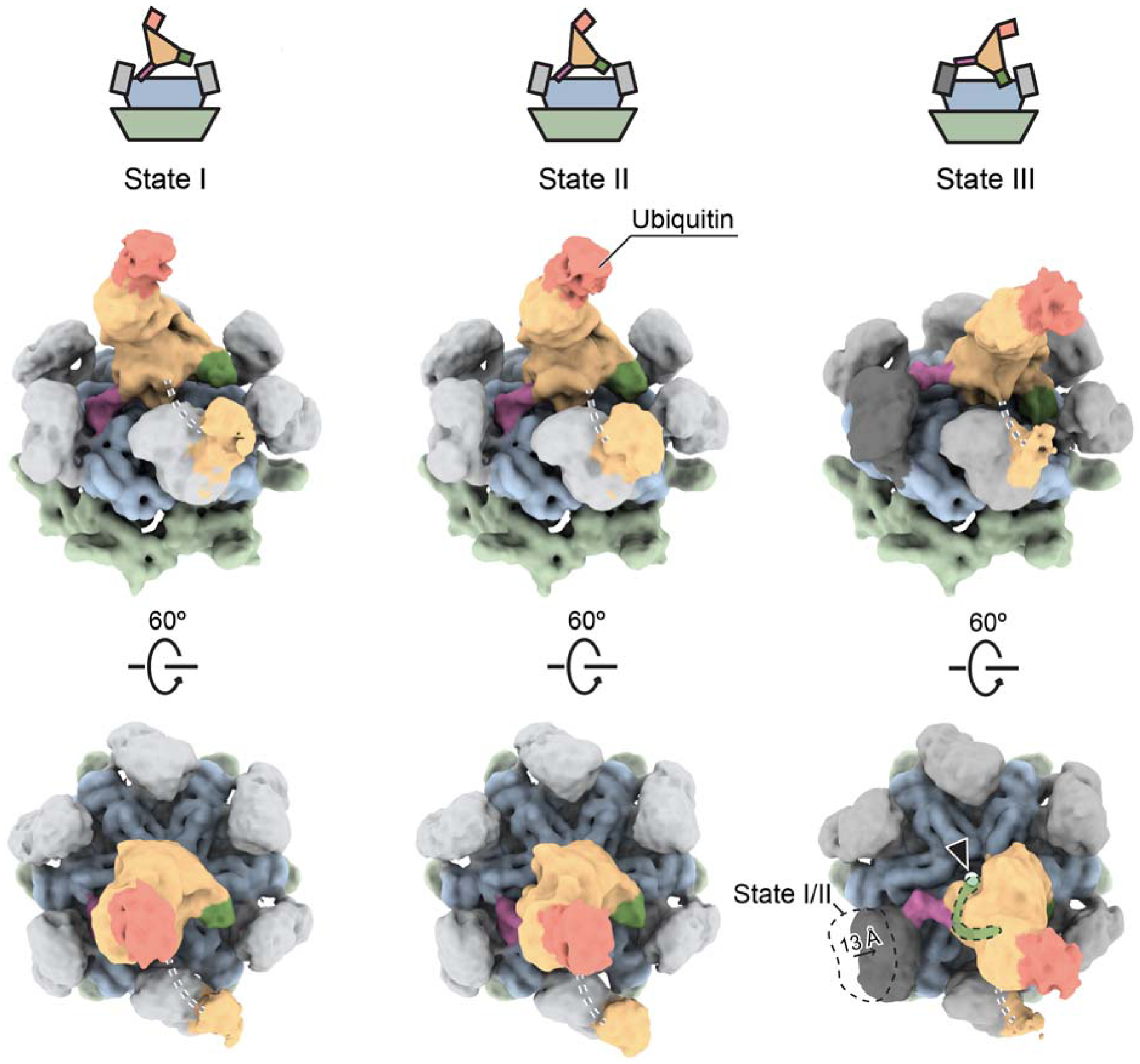
Three conformational states of human p97 in complex with Npl4/Ufd1 and polyubiquitinated Ub-Eos. Single-particle cryo-EM maps (unsharpened) of human p97 in complex with Npl4/Ufd1 and polyubiquitinated Ub-Eos. Three conformational states were resolved using 3D classification. Each state is in a similar conformation to one of the states in the complex of p97 and Npl4/Ufd1 without the substrate (compare to Fig. 1b). One ubiquitin molecule (pink) was observed to bind to the top of Npl4. One of the N domains in State III (dark gray) shifted and interacted with ZF1 (zoomed in Supplementary Fig. 4b).The color scheme used for the other domains is the same as that used in Fig. 1. The position of Npl4 relative to p97 is illustrated in the cartoons. Unresolved linkers between UBXL and ZF1 are represented by dotted lines. The central pore of the D1 ring in State III is highlighted by an arrowhead. The conserved groove of Npl4 that was found to bind an unfolded ubiquitin in the yeast complex structure^5^ was marked by a green band in State III.

### Seesaw motion of Npl4 is important for the unfolding activity of human p97

To further investigate the role of the seesaw motion of Npl4, we determined the structure of human p97 in complex with Npl4/Ufd1 and a K48-linked polyubiquitinated substrate Ub-Eos, which has two tandem ubiquitin molecules fused to the N-terminus of the fluorescent protein mEos3.2, using single-particle cryo-EM (**Supplementary Fig. 3**). To stabilize the complex, we introduced two mutations into p97, A232E and E578Q. It has been reported that A232E, a mutation found in patients with neurodegenerative diseases, has increased affinity to Npl4/Ufd1^16,17^. E578Q, a mutation in the Walker B motif of the D2 domain, inhibits ATP hydrolysis in the D2 ring. No additional nucleotides were supplied during the purification. Again, three major conformational states of the p97-Npl4/Ufd1 complex with substrate engagement were resolved (**Fig. 2**). In substrate-engaged States I and II, the conformations are essentially the same as those observed from States I and II without the substrate. The only change was the observation of density corresponding to a single ubiquitin on the very top of Npl4. In substrate-engaged State III, in addition to the ubiquitin density, we also found a displacement of one N domain that was “dragged” by the raised ZF1 motif, which is approximately 13 Å (**Fig. 2**). The density corresponding to the polyubiquitinated Ub-Eos is only visible at a lower threshold and is not resolvable (**Supplementary Fig. 4a**). Recently, the cryo-EM structure of yeast Cdc48 in complex with Npl4/Ufd1 and K48-linked polyubiquitinated Eos showed that an unfolded ubiquitin moiety binds in the groove of Npl4 and extends all the way into the pore of Cdc48 in the absence of ATP hydrolysis^5^. By contrast, none of the states showed any density in the groove of Npl4 (**Fig. 2**), suggesting that the resolved structures are prior to ubiquitin unfolding and the polyubiquitinated substrate translocating. Together, the main conformational change caused by the binding to the ubiquitinated substrate is the clear interaction between ZF1 and the N domain of p97 in State III. (**Fig. 2 and Supplementary Fig. 4b**). Since the resolutions of the maps do not allow model building at the atomic level (**Supplementary Fig. 3b**), we docked the crystal structure of the N domain^18^ and the homology model of Npl4 into the density as rigid bodies, which showed that R113 in the N domain is likely involved in the interaction (**Supplementary Fig. 4b**). To test if the interaction is relevant for the function of p97, we mutated the residue R113 to an alanine in the N domains of wild-type p97 and performed a previously established fluorescence-based substrate unfolding assay (**Fig. 3a**)^16^. In this assay, the decrease in the fluorescence signal corresponds to the unfolding of the polyubiquitinated Ub-Eos substrate (**Fig. 3b**). As shown in the results, the unfolding activity of R113A was greatly reduced to a similar level as the A232E/E578Q double mutant, suggesting that the interaction is indeed important for the unfolding activity of human p97 (**Supplementary Fig. 4c and d**). In summary, our structures revealed unexpected conformational changes of Npl4 before ATP hydrolysis and translocation, which were not observed in the fungal homolog^5,6^. The unfolding of ubiquitin did not occur in our complex, suggesting a mechanistic difference between human 97 and yeast Cdc48 (see discussion). Most interestingly, our structures uncovered an important role of the zinc finger motifs in Npl4, one of which had been recently identified as the binding site of the disulfiram derivative, CuET^8^. Therefore, we conducted further investigations on how CuET might affect the function of human p97.

**Figure 3:**
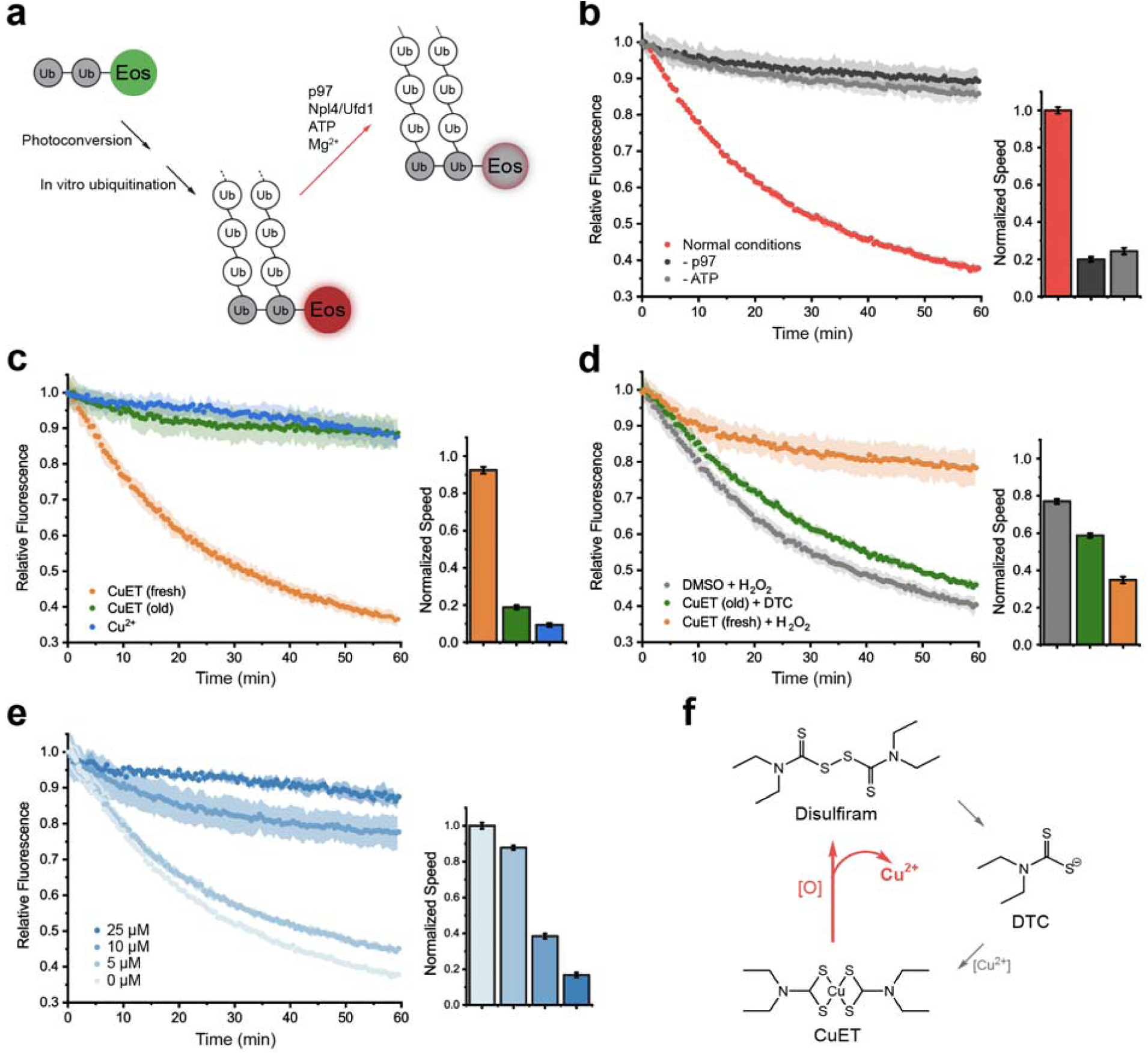
Copper released from the disulfiram derivative CuET under oxidative conditions inhibits the unfolding activity of p97. **a**, A diagram showing the established substrate unfolding assay of p97. The unfolding of ubiquitinated Eos corresponds to a decrease in the fluorescence signal (red arrow). **b**, The unfolding activity of wild-type p97 in the presence of Npl4/Ufd1, 5 mM ATP, 20 mM MgCl_2_ and ubiquitinated Eos (normal conditions). The relative fluorescence signal (starting at the time point zero) was monitored for 60 minutes. The initial velocity of the reaction was linearly fitted using the data points from the first 10 minutes and plotted in the bar graph that was normalized against the normal condition. **c**, The unfolding activity and initial velocity of p97 in the presence of fresh or old CuET (50 μM) or cupric ions (25 μM). **d**, The unfolding activity and initial velocity of wild-type p97 in the presence of fresh or old CuET (50 μM) and an oxidative (H_2_O_2_, 100 μM) or a reducing (DTC, 100 μM) reagent. **e**, The unfolding activity of p97 is inhibited by cupric ions in a concentration-dependent manner. **f**, The cycle of disulfiram derivatives. CuET can release cupric ions under oxidative conditions (red arrow). The error bands and bars in panels **b-e** represent the standard deviation from triplicate experiments.

**Figure 4:**
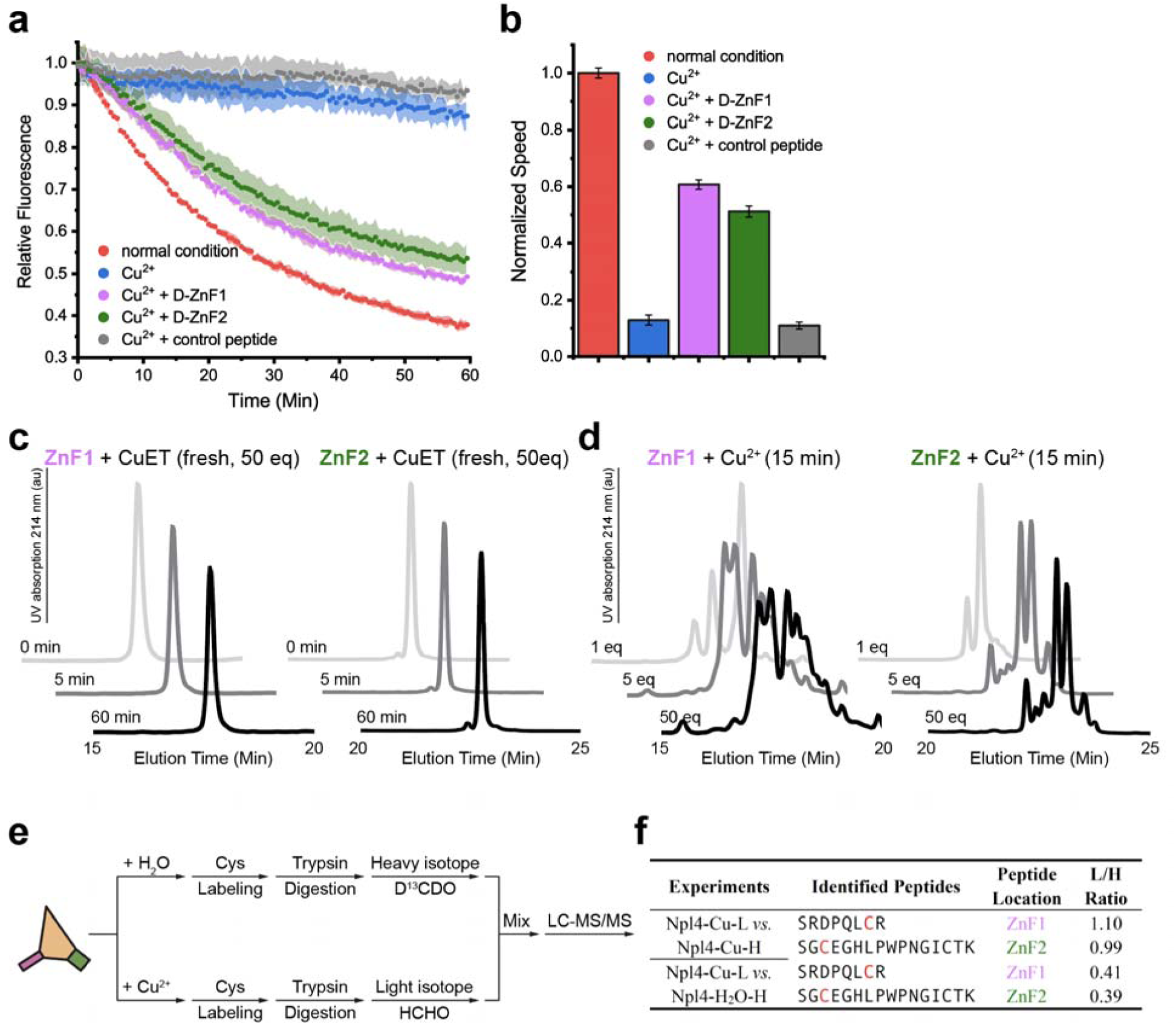
Copper interacts with the zinc finger motifs of Npl4. **a**, Zinc finger motifs of Npl4 rescue the unfolding activity of p97 in the presence of cupric ions. **b**, The initial velocity corresponds to panel a. The error bands and error bars in panels a and b represent the standard deviation from three experimental repeats. **c**, HPLC profiles of synthesized ZF1 and ZF2 peptides mixed with fresh CuET (at 1:50 molar ratio) over 60 minutes. **d**, HPLC profiles of synthesized ZF1 and ZF2 peptides mixed with cupric chloride at different ratios for 15 minutes. **e**, A diagram showing the experimental steps of quantitative mass spectrometry using light and heavy formaldehyde to probe the interactions between cupric ions and the cysteine residues in the zinc finger motifs of Npl4. Cupric ions oxidize the cysteine residues of the zinc finger motifs and decrease the labeling percentage. **f**, Results from the quantitative mass spectrometry (panel e). Labeled cysteine residues are highlighted in red. The first experiment was used as a control, in which both light and heavy labeling were treated with cupric ions, and the expected L/H ratio was 1.

### Disulfiram derivative CuET inhibits the function of p97 through copper release

Recently, the metabolic derivative of the anti-alcohol abuse drug disulfiram, CuET (**Supplementary Fig. 5a**), has been reported to bind to ZF1 of Npl4^8^. Since the zinc finger motifs in Npl4 play critical roles in different conformational states, we tested whether CuET affects the unfolding activity of p97. We synthesized CuET using diethyldithiocarbamate (DTC) and cupric chloride. At room temperature, CuET forms brown-colored crystals and is soluble in dimethyl sulfoxide (DMSO) (**Supplementary Fig. 5b**). Unexpectedly, we found that only old CuET solution (left at 4 °C for more than 7 days) could inhibit the unfolding activity of p97 (**Fig. 3c, green curve**); freshly dissolved CuET showed minimal effect (**Fig. 3c, orange curve**). It is known that disulfide bonds may form in the presence of DMSO^19^. Therefore, we tested if cupric ions were present in the old CuET solution due to the oxidation of the dithiocarbamate group (**Supplementary Fig. 5c**). Indeed, as we expected, cupric ions were released from the molecule, and oxidative reagents such as hydrogen peroxide can accelerate the reaction, leading to a dramatic color change in the CuET solution (**Supplementary Fig. 5d**). The reaction depends on the concentration of the oxidant. With a small amount of oxidant, it can last for days (**Supplementary Fig. 5d**). We further tested if the released cupric ions could actually inhibit the unfolding activity of p97. Indeed, cupric ions inhibited the activity in a concentration-dependent manner (**Fig. 3c and e)**. The release of cupric ions suggested that there is a cycle of disulfiram, DTC, and CuET (**Fig. 3f**). To further test the existence of a cycle, we introduced additional DTC into the unfolding assay in the presence of old CuET solution. As expected, DTC can rescue the unfolding activity of p97 by chelating the released cupric ions (**Fig. 3d, green curve**). We also introduced hydrogen peroxide into the unfolding assay in the presence of fresh CuET solution and found that hydrogen peroxide can inhibit the activity by triggering the release of cupric ions (**Fig. 3d, orange curve**). Furthermore, we measured the unfolding activity of p97 in the presence of other metal ions at the same concentration (25 μM). Interestingly, cupric ions appeared to be the most potent inhibitor in vitro (**Supplementary Fig. 5e**).

**Figure 5:**
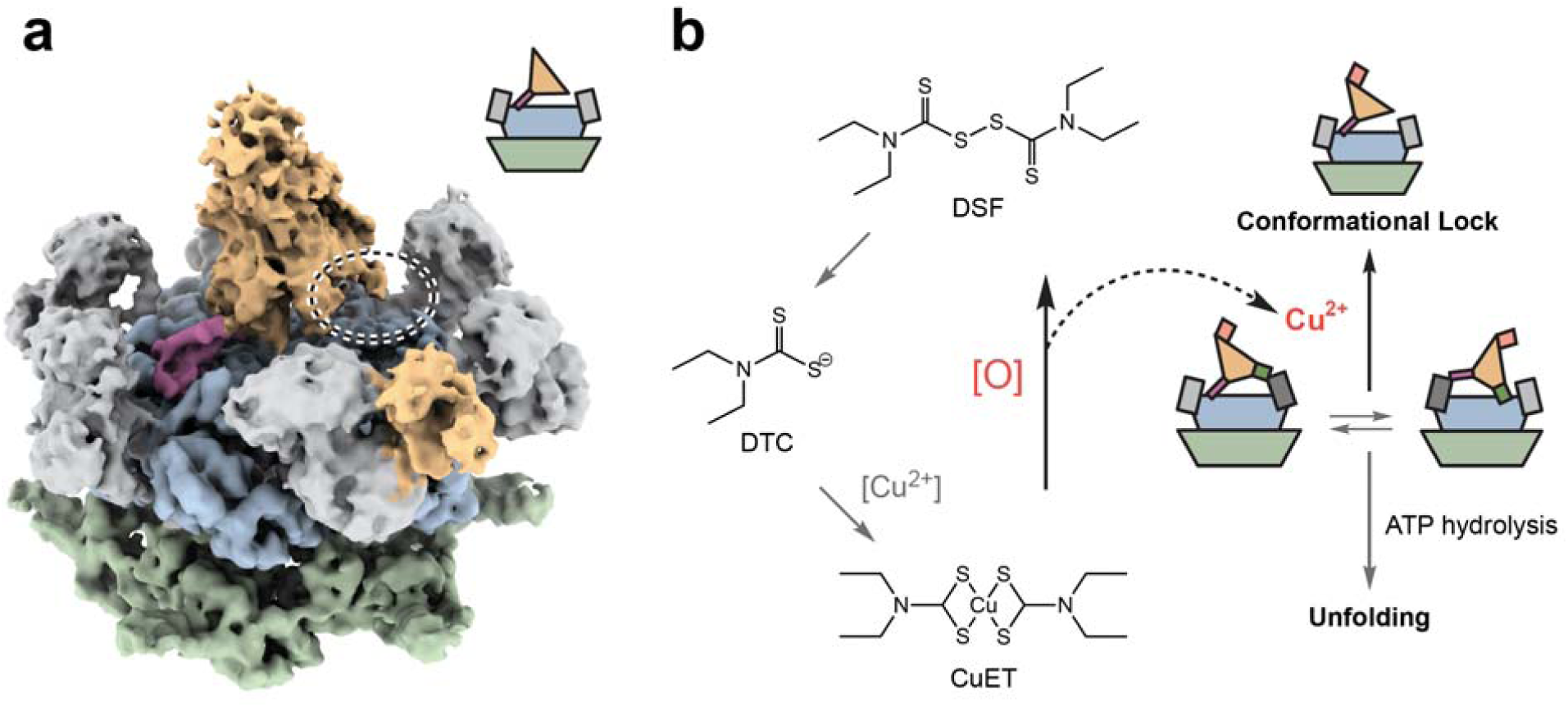
Conformational lock induced by copper released from CuET. **a**, Single-particle cryo-EM map (unsharpened) of human p97 in complex with Npl4/Ufd1 in the presence of cupric ions (100 μM). The color scheme is the same as that used in Fig. 1. In contrast to the complex without cupric ions, only one conformational state was resolved. Note that ZF2 was not resolved in the cryo-EM map (the circle highlights this area). **b**, A summary of the findings in this work. CuET releases cupric ions under oxidative conditions, inhibiting the unfolding activity of p97 through conformational locking.

To test the relevance of our findings in live cells, we first determined the dose response curves of CuET and the oxidant, tert-butyl hydroperoxide (tBHP), for cultured HeLa and A549 cells (**Supplementary Fig. 6a-c**). Oxidative stress was induced by treating the cells with tBHP for 6 hours. CuET (250 nM) was introduced after the treatment, and the cells were incubated for another 20 hours; then, an MTT assay was used to monitor cell survival (**Supplementary Fig. 6d**). Compared to control cells that had not been subjected to oxidative stress, the cell survival was significantly lower (p < 0.01) for both CuET-treated HeLa and A549 cells (**Supplementary Fig. 6e and f**). Our findings suggested that CuET is more toxic to the two cancer cell lines under oxidative conditions. Since the cells are not permeable to cupric ions, the toxicity is likely due to the release of cupric ions in the cytosol from CuET, which can cross the cell membrane (see discussion).

### Cupric ions interact with the zinc finger motifs of Npl4

It is known that copper can interact with cysteine residues in zinc finger motifs^20^. Npl4 has three zinc finger motifs: ZF1 and ZF2 bind to the D1 ring and the N domains of p97; NZF binds to the polyubiquitin chains. We therefore tested if cupric ions affected the complex formation of p97, Npl4/Ufd1, and polyubiquitinated Ub-Eos. We formed the complex in the presence of cupric chloride, followed by size-exclusion chromatography and SDS-PAGE. The results showed that cupric ions do not affect the binding between Npl4 and p97 (**Supplementary Fig. 7a**), nor do they affect the binding between Npl4 and the polyubiquitinated Ub-Eos (**Supplementary Fig. 7b**). To further quantify the interaction between Npl4 and the copper compounds, we performed a series of isothermal titration calorimetry (ITC) experiments (**Supplementary Fig. 8**). The results showed that freshly dissolved CuET does not interact with Npl4 (**Supplementary Fig. 8c**). Instead, cupric glycinate (used as a replacement for cupric ions) binds to Npl4 at an apparent dissociation constant (Kd^app^) of 2.8 ± 1.1 μM (**Supplementary Fig. 8d**). By comparison, the binding between cupric glycinate and p97 is much weaker, with a Kd^app^ of 69 ± 18 μM (**Supplementary Fig. 8e**). We then tested the binding between cupric glycinate and Npl4 bearing zinc finger motif mutants (**Supplementary Fig. 8a**). With a ZF1 mutation, the binding constant is 7.8 ± 2.0 μM; with ZF2 mutation, the binding constant is 8.7 ± 1.8 μM; and with both ZF1 and ZF2 mutated, the binding constant is 20 ± 6.5 μM (**Supplementary Fig. 8f-h**). Taken together, the results suggested that cupric ions interact with Npl4 primarily through ZF1 and ZF2.

To test if the interaction between cupric ions and zinc finger motifs of Npl4 inhibits the unfolding activity of p97, we performed a substrate unfolding assay in the presence of cupric ions and used synthesized zinc finger motifs with D-amino acids (**Supplementary Fig. 8a and b)** to rescue the activity (**Fig. 4a and b**). The rationale is that the synthesized D-zinc finger motifs can interact with cupric ions in the same way as the normal L-zinc finger motifs but cannot interact with p97 due to inverted chirality; hence, the seesaw motion of Npl4 will not be disturbed. Therefore, if cupric ions inhibit the unfolding activity of p97 by interacting with the zinc finger motifs of Npl4, the synthesized D-zinc finger motifs should be able to rescue the reactivity. Indeed, both D-ZF1 and D-ZF2 could rescue the unfolding activity of p97, whereas the control D-peptide from Npl4 (**Supplementary Fig. 8a and b**) could not (**Fig. 4a and b**).

It has been reported that copper ions could disrupt the zinc finger structure by mediating the oxidation of cysteine residues^20^. Therefore, we further analyzed the reaction between the copper compounds and synthesized zinc finger motifs using high-performance liquid chromatography (HPLC). The results showed that after 60 minutes of incubation at a 50:1 (Cu^2+^ to peptide) molar ratio, the freshly dissolved CuET did not react with either zinc finger motif (**Fig. 4c**). By contrast, cupric ions could react with both peptides in a concentration-dependent manner after 15 minutes of incubation, suggesting that the structures of both zinc finger motifs can be disrupted by cupric ions (**Fig. 4d**). To analyze how the zinc finger motifs in Npl4 react with cupric ions, we performed an iodoacetyl-modified probe-based quantitative mass spectrometry analysis for the cysteine residues in the zinc finger motifs of Npl4. We first incubated purified Npl4 with or without cupric chloride for 30 minutes at room temperature, followed by the following parallel treatment: cysteine labeling, trypsin digestion, and isotope labeling by light and heavy formaldehyde. The reactants were mixed for quantitative liquid chromatography with tandem mass spectrometry (LC-MS/MS) analyses (**Fig. 4e**). The results showed that the cysteine residues in both zinc finger motifs were much less labeled (light/heavy (L/H) ratio less than 1) after incubation with cupric ions, suggesting that those cysteine residues had been oxidized and were not subjected to the labeling reaction (**Fig. 4f**).

### Conformational locking upon copper release

Finally, we determined the single-particle cryo-EM structure of the p97-Npl4/Ufd1 complex in the presence of cupric ions (**Fig. 5a**). With extensive 3D classifications, we only observed one dominating conformation that is similar to State II of the same complex without cupric ions (**Fig. 1b and Supplementary Fig. 9**). The final reconstruction reached 3.5 Å, the highest resolution in this study, but the Npl4 part of the cryo-EM map is less well resolved than that of the other regions. Specifically, the ZF2 motif was not resolved in the cryo-EM map, suggesting that under the experimental conditions, ZF2 may have been disrupted by cupric ions and could not bind to the D1 ring of p97. Therefore, we did not obtain a conformation similar to State III of the same complex without cupric ions. Since the seesaw motion of Npl4 is important for the unfolding activity of human p97, the lack of other states suggested a conformational lock, leading to the inhibition of activity.

## DISCUSSION

In this study, we focused on structural and functional analyses of human p97 and its cofactor Npl4/Ufd1. We discovered three major conformational states of the p97-Npl4/Ufd1 complex both in the presence and absence of the polyubiquitinated substrate. Our structures of human p97 complexes suggest a seesaw motion of Npl4 on top of the D1 ring. We further showed that the seesaw motion is essential for unfolding the substrate. Note that one of the states exposes the central pore of the D1 ring. This dynamic movement leads to a model: the seesaw motion of Npl4 may facilitate the initial unfolding of ubiquitin by exposing the pore of the D1 ring. The partially unfolded or destabilized ubiquitin is then captured by Npl4 or directly inserted into the pore, which may or may not lead to a structure similar to that observed for yeast Cdc48. The latter possibility suggests a different mechanism between yeast and humans, which cannot be ruled out at this stage. There are still missing pieces in this model, and further structural studies are required to elucidate the details.

We also found that at least one zinc finger motif interacts with the N domains of p97 (**Supplementary Fig. 4b**). It is known that N domains change conformations upon ATP hydrolysis^21,22^. Therefore, the seesaw motion we observed here may be coupled with ATP hydrolysis that occurs in the D1 and D2 rings through the interactions between the zinc finger motifs and the N domains. In such a case, ATP hydrolysis would be the driving force for the seesaw motion of Npl4. What we captured without triggering ATP hydrolysis are three intermediate conformations of the motion. Note that this does not contradict the model of translocation initiation that is proposed above, as initiation may require ATP hydrolysis as well.

We showed that CuET could disrupt the seesaw motion of Npl4 through the release of copper ions under oxidative conditions (**Fig. 5b**). This is likely a mechanism linking disulfiram to p97 inhibition. Disulfiram has long been known to have anticancer activity^23^. Recently, the metabolic derivative of disulfiram, CuET, has been found to target cancer though Npl4^8,24^. Our biochemical and structural data suggested that cupric ions, not CuET, interact with the zinc finger motifs of Npl4; as a result, cupric ions inhibit the unfolding activity of p97 through a conformational lock. We discovered that cupric ions are released by CuET under oxidative conditions. The velocity of the reaction depends on the concentration of the oxidant and can last for days (**Supplementary Fig. 5d**). Such conditions may exist in cancer cells and tissues that are often under oxidative stress^25^. Therefore, we propose that instead of directly targeting Npl4, CuET acts as a shuttle for cupric ions to cross the cell membrane, similar to the mechanism reported for other copper chelators^26,27^. The development of copper-based anticancer drugs has been very popular^28^, but the transport of copper into cells is strictly regulated^29^. CuET can bypass the copper transporter system and freely penetrate cells since the intracellular copper concentration increased rapidly when the cells were treated with disulfiram in medium containing cupric chloride, and the intracellular copper uptake could be blocked by coincubation with bathocuproinedisulfonic acid, a nonmembrane-permeable copper chelator^30^. Once in the cytosol, CuET releases cupric ions depending on the cellular redox status, which then impair the function of essential cellular machines such as p97. Although there are many cellular processes that may be affected by cupric ions^31,32^, p97 may be one of the major targets, as it is very abundant and plays a key role in cellular proteostasis. Further investigations are required to elucidate the relationship between cytosolic cupric ions and the function of p97 in a cellular environment.

## ACKNOWLEGEMENT

Funding for this work was, in part, provided by the Catalyst Award from the Chicago Biomedical Consortium. This work was supported by Chicago Biomedical Consortium Catalyst Award C-086 to M.Z. We thank Zhiheng Yu and Doreen Matthies at the HHMI Janelia CryoEM Facility for the help in cryo-EM data collection. This research was, in part, supported by the National Cancer Institute’s National Cryo-EM Facility at the Frederick National Laboratory for Cancer Research under contract HSSN261200800001E.

## AUTHOR CONTRIBUTIONS

M.P., L.L. and M.Z. designed all the experiments and interpreted the results. M.P., Q.Z., Y.Y., and H.A. cloned, expressed and purified all the complexes and carried out related biochemical characterizations including the substrate unfolding assay and the ITC experiments. M.P. and M.Z. performed cryo-EM data collection and processing. Y.Y. synthesized the disulfiram derivative CuET. Y.X. performed the cell survival assay. Q.Z., Z.X. and C.W. performed the iodoacetyl modified probe based LC-MS/MS experiments. M.Z., M.P., and Q.Z. wrote the paper. M.Z. and L.L. supervised the project.

## CORRESPOINDING AUTHOR

Correspondence to Minglei Zhao and Lei Liu.

## COMPETING INTERESTS

The authors declare no competing interests.

## DATA AVAILABILITY

cryo-EM maps have been deposited in the Electron Microscopy Data Bank (EMDB) under the accession codes EMDB-21824 (p97-Npl4/Ufd1, State I), EMDB-21825 (p97-Npl4/Ufd1, State II), EMDB-21826 (p97-Npl4/Ufd1, State III), EMDB-21827 (p97-Npl4/Ufd1-UbEos, State I), EMDB-21828 (p97-Npl4/Ufd1-UbEos, State II), EMDB-21829 (p97-Npl4/Ufd1-UbEos, State III), and EMDB-21830 (p97-Npl4/Ufd1 in the presence of cupric ion).

## METHODS

### Protein expression and purification

Full-length human p97 including mutant A232E/E578Q, Ufd1, Npl4, as well as wild-type and mutant Npl4 fragments (residues 129–580) were overexpressed in *E. coli* BL21-DE3-RIPL cells. p97 was cloned into pET-47b vector with an N-terminal cleavable His-tag. Two plasmids containing full-length untagged Npl4 and C-terminal His-tagged Ufd1 were purchased from Addgene. Npl4 fragments were expressed in pET-28a vector with an N-terminal His-SUMO tag. Purification protocols were similar to those published before^33^. Briefly, *E. coli* cells were grown in LB with auto-induction supplement for 16 h at 30 °C. Cells were pelleted at 5,000 g and resuspended in Lysis Buffer (50 mM Tris, pH 8.0, 300 mM KCl, 20 mM imidazole, 1 mM DTT and 1 mM MgCl_2_). To purify the Ufd1-Npl4 complex, the resuspended cells of respective construct were mixed at this step. 1mM PMSF was added to the suspensions, and cells were lysed by sonication. Lysates were cleared by centrifugation for 30 min at 16500 rpm. then the supernatants were flowed through a Ni–NTA gravity column twice at 4 °C. Beads were washed with Wash Buffer (50 mM Tris, pH 8, 150 mM NaCl, 20 mM imidazole and 1 mM MgCl_2_). The His-tagged proteins were eluted in Elution Buffer (50 mM Tris, pH 8.0, 300 mM NaCl, and 400 mM imidazole). For p97 and Npl4 fragments, HRV3C protease and SUMO protease (Ulp1p) were added to the eluate, respectively, and dialyzed in Buffer A (25 mM Tris, pH 8.0, 150 mM NaCl, 1 mM MgCl_2_, and 0.5 mM Tris(2-carboxyethyl)phosphine (TCEP)). Ufd1-Npl4 complex were directly dialyzed in Buffer A. Ufd1-Npl4 complex and Npl4 fragments were further purified by Superdex 200 size-exclusion column (GE Healthcare) equilibrated in Buffer A. p97 was purified by Superose 6 column (GE Healthcare).

### Preparation of poly-ubiquitinated Ub-Eos

Ub-Eos was constructed by fusing two tandem ubiquitin to the N-terminus of fluorescent protein mEos3.2. The poly-ubiquitinated Ub-Eos was prepared similar to previously described^16^. Briefly, ubiquitination reaction was carried out with final concentrations of 20 μM Ub-Eos, 1 μM E1, 20 μM gp78RING-Ube2g2, and 500 μM ubiquitin in 20 mM Hepes, pH 7.4, 150mM KCl, 10 mM ATP, and 10 mM MgCl_2_ at 37 °C overnight. To purify ubiquitinated Eos3.2 from free ubiquitin chains, the reaction mixture was incubated with Ni-NTA resin, eluted with 300 mM imidazole, and run over a Superdex 200 size-exclusion column in Buffer A. Fractions with long ubiquitin chain (over 10 Ubs) were pooled and concentrated with centrifugal filter units, followed by flash-frozen.

### p97 Complex formation

To form p97-Ufd1-Npl4 complex, Ufd1-Npl4 was added at a three folds molar excess to p97 hexamer, and proteins were incubated with 1 mM ATP-γS for 60 min before gel filtration. After gel filtration, all proteins were flash frozen in liquid nitrogen. For p97-Ufd1-Npl4-Poly-Ub Eos complex, a mutant p97 bearing A232E and E578Q mutations was used instead of wild-type p97. To form the ternary complex, p97 (A232E/ E578Q)-Npl4/Ufd1 complex was first prepared as described above, then poly-ubiquitinated Ub-Eos was added at a two-fold molar excess to p97 (A232E/E578Q)-Npl4/Ufd1 complex. After gel filtration, all proteins were flash frozen in liquid nitrogen.

### Specimen preparation for single-particle cryoEM

For p97-Npl4/Ufd1 complex, before preparing grids for cryo-EM, the complex was concentrated to ∼20 mg/ml, and incubated with 5 mM ATPγS (90% pure, Sigma-Aldrich) for 30 min at room temperature. For p97 (A232E/E578Q)-Npl4/Ufd1-UbEos complex, the sample was concentrated to 20mg/ml without adding additional nucleotide. For p97-Npl4/Ufd1 complex in the presence of cupric ion, the complex was concentrated to ∼20 mg/ml, and incubated with 5 mM ATPγS (90% pure, Sigma-Aldrich) and 100 μM cupric ion for 30 min at room temperature. To relief the preferred orientations, IGEPAL CA-630 (Sigma-Aldrich) was added to the samples to a final concentration of 0.05% immediately before grid freezing. The freezing was performed using a Vitrobot mark IV (Thermo Fisher). Samples (3.5 μL) were applied to a non-glow-discharged Quantifoil Au 1.2/1.3 grid. The grid was blotted for one second and then plunge frozen in liquid ethane.

### Data collection for single-particle cryoEM

Optimized frozen grids were shipped to National Cryo-Electron Microscopy Facility for data collection. All datasets were acquired as movie stacks with a Titan Krios electron microscope operating at 300 kV, equipped with either a Gatan K2 Summit or K3 direct detector camera. A single stack typically consists of 40 frames with a total exposure around 50 electrons/Å^**2**^. The defocus range was set at -1.0 to -2.5 μm. See **Supplementary Table 1** for the details.

### Image Processing

Movie stacks were subjected to beam-induced motion correction using MotionCor2^34^. CTF parameters for each micrograph were determined by CTFFIND4^35^. The following particle picking, two- and three-dimensional classifications, and three-dimensional refinement were performed in RELION-3^36^. Briefly, false-positive particles or particles classified in poorly defined classes were discarded after 2D classification. The initial 3D classification was performed on a binned dataset with the previously reported p97 structures as the reference model^22^. The detailed data processing flows are shown in Supplementary Figs. 1, 3, and 9. To make sure that the 3D classification did not miss any major conformations, parallel runs were performed asking for different number of classes, which ended up with the same number of major classes. Data processing statistics are summarized in Supplementary Table 1. Reported resolutions are based on Fourier shell correlation (FSC) using the FSC=0.143 criterion. Local resolution was determined using ResMap^37^ with half-reconstructions as input maps.

### Model Building, Refinement, and Validation

Model building was based on the existing crystal structures of human p97^12^ (PDB code: 3CF3). A homology model of human Npl4 was built using SWISS-MODEL^38^ based on the crystal structure of thermophilic fungi Npl4^6^ (PDB code: 6CDD). The existing p97 model and the homology model were first docked into the cryoEM maps as rigid bodies using UCSF Chimera^39^. The p97 part was then manually adjusted residue-by-residue to fit the density using COOT^40^, and was subjected to global refinement and minimization in real space using the real space refine module in Phenix^41^. The Npl4 part was kept as a rigid body during the process.

### Cys reactivity validation through iodoacetyl modified probe based LC-MS/MS

The quantitative chemical proteomics was performed as recently described^42^. 200 μL Npl4 (0.11 mg/mL in 25 mM Hepes, pH 7.6, 150 mM NaCl,) was directly incubated with 1μL CuCl_2_ (2mM) or 1 μL ddH_2_O for 30 minutes at 25 °C, respectively. After the treatment with or without copper, iodoacetamide (IA probe) was added to a final concentration of 100uM, and incubated for 60 minutes at 25 °C. Then, 10 μL DTT (200mM) was added to each sample at 37 °C for 30 min to quench the reaction. Each solution was diluted with 1ml TEAB to a final urea concentration of 2M. 10 μL mass spectrometry grade trypsin (0.5 μg/μL, ratio of samples: trypsin = 1 : 40 (w/w), Promega, V5280) was used to digest protein samples overnight at 37 °C. 8 µL of 4% (v/v) ‘light’ (Sigma Aldrich, F1635) or ‘heavy’ formaldehyde-13C, d2 (Sigma Aldrich, 596388) was added to the reaction with or without the copper treatment, respectively. 8 µL of sodium cyanoborohydride (0.6 M) were added and the reaction was incubated at 25 °C for 1 hour before quenching with 32 µL of 1% ammonia followed by 16 µL of 5% formic acid. The corresponding ‘light’ and ‘heavy’ sample were combined and centrifuged (1,400 g, 2 min). Mixed peptides were dried and stored at -20 °C until LC-MS/MS analysis.

LC-MS/MS data was analyzed by ProLuCID^43^ with static modification of cysteine (+57.0215 Da) and variable oxidation of methionine (+15.9949 Da). The isotopic modifications (+28.0313 and +34.0631 Da for light and heavy labeling respectively) were set as static modifications on the N-terminal of a peptide and lysine residues. Additional 357.17223 Da of IA probe was set as variable modifications on cysteines. The searching results were filtered by DTASelect^44^ and peptides were also restricted to fully tryptic with a defined peptide false positive rate of 1%. The ratios (L/H) of reductive dimethylation were quantified by the CIMAGE software as described before^45^.

### Substrate unfolding assay

The polyubiquitinated, photo-converted Ub-Eos was prepared as described^17^. Experiments were carried out in Reaction Buffer (20 mM Hepes, pH 7.4, 150 mM KCl, 20 mM MgCl2, 1 mg/mL BSA) supplemented with an ATP regeneration mixture (5 mM ATP, 30 mM creatine phosphate, and 50 μg/mL creatine phosphokinase). Proteins were pre-incubated in a 96-well plate (Fisherbrand FB012931) for 10 minutes at 37 °C before adding the ATP regeneration mixture to initiate the reaction. Final concentrations of the reactants were 20 nM substrate, 400 nM p97 in hexamer, and 300 nM Ufd1-Npl4. Fluorescence signal was monitored using a TECAN safire2 plate reader at 540 nm excitation and 580 nm emission wavelength and 30 s intervals for 60 min. Each reaction was repeated three times. Background fluorescence was measured by mixing the same amount of substrate with 6 M guanidine-HCl and was subtracted from the average of the experimental groups. Normalized fluorescence and the initial velocity of the reaction (first 20 data points corresponding to 10 min of reaction) were plotted and fitted using OriginPro (OriginLab).

### Isothermal titration calorimetry

Isothermal titration calorimetry measurements were carried out on an ITC200 Microcalorimeter (GE Healthcare). p97 and Npl4 fragments were dialyzed in a buffer containing 25 mM HEPES (pH 7.4) and 150 mM NaCl. Sufficient amount of 1 mM Cu(Gly)_2_ in the injection syringe was titrated to the sample cell containing 0.02 mM of p97 or Npl4 fragments to achieve a complete binding isotherm. All binding experiments were performed at constant temperature of 25 °C. A total of 20 injections of 2.0 μL were dispensed with a 5-second addition time and a spacing of 120 seconds. All experiments were repeated at least three times. Data were analyzed and the titration curves were fitted using MicroCal Origin software assuming a single binding site mode.

### Peptide synthesis

All peptides used in this work were synthesized using standard Fmoc SPPS protocols under microwave conditions (CEM Liberty Blue)^46^. Rink Amide AM resin was first swelled in DMF for 10 min. Every coupling cycle was executed programmatically. In general, the deprotected condition is 10% piperidine in DMF with 0.1 M Oxyma (1 min at 90 °C) and the amino acid coupled condition is 4-fold of 0.2 M Fmoc-protected amino acid, 1.0 M DIC, and 1.0 M Oxyma in DMF (10 min at 50 °C for His and Cys, 90 °C for other residues). After the completion of SPPS, the peptide-resin was transferred into customized sand core funnel and treated with the cleavage cocktail (TFA/H2O/thioanisole/EDT 87.5/5/5/2.5, v/v/v/v) for 2 h at room temperature. Crude peptides were precipitated with cold diethyl ether and dissolved in water (containing 0.1% TFA) mixed with acetonitrile (containing 0.1% TFA) and purified by semi-preparative RP-HPLC.

### MTT Assay for CuET and copper toxicity

The MTT assay to evaluate CuET toxicity under oxidative conditions in cancer cells was modified from previous protocols^47^. Briefly, cells were plated on 96-well tissue culture plates (Fisher, FB012931) at 5,000 cell per well and grown at 5% CO2 at 37 °C. The next day, oxidative stress was achieved by treating the cells with growth medium containing 0, 25, or 50 μM TBH70X (Sigma, 458139). After 6 hours, the TBH70X-containing medium was removed and replaced with medium containing 0, 0.1 or 0.25 μM CuET. The DMSO control contained 100 μl culture medium with 0.05 μl DMSO (Fisher, BP231100). After 18 hours of CuET treatment, 20 μL of 5 mg/mL MTT (Sigma, M2128) was added to each well and the cells were further incubated for 3.5 hours in 37 °C incubator. To dissolve the formazan crystals, the culture medium was carefully removed and 150 μL of MTT solvent (4mM HCl, 0.1% NP40 in isopropanol) was added to each well with 15 min shaking at room temperature. The absorbance was measured at 590nm with a reference wavelength of 620nm with a SAFIRE II plate reader (Tecan, Männedorf, Switzerland). Each experimental group contained 4 biological replicates with the error bar indicating standard deviation. The cell survival rate was calculated by the following equation: cell survival rate (%) = (absorbance of experimental group/absorbance of control group) × 100%.

**Supplementary Table 1:** Statistics of cryo-EM data collection and processing.

|  | p97-Npl4/Ufd1 |  |  | p97-Npl4/Ufd1-UbEos |  |  | p97-Npl4/Ufd1-Cu<sup>2+</sup> |
| --- | --- | --- | --- | --- | --- | --- | --- |
| Magnification | 105,000 |  |  | 64,000 |  |  | 81,000 |
| Voltage (kV) | 300 |  |  | 300 |  |  | 300 |
| Detector | K2-Summit |  |  | K3 |  |  | K2-Summit |
| Electron exposure (e<sup>-</sup>/Å<sup>2</sup>) | 50 |  |  | 50 |  |  | 50 |
| Defocus range (μm) | -1.0 to -2.5 |  |  | -1.0 to -2.5 |  |  | -1.0 to -2.5 |
| Pixel size (Å) | 0.66 |  |  | 0.68 |  |  | 1.01 |
| Symmetry imposed | C1 |  |  | C1 |  |  | C1 |
| Initial particle images (no.) | 370,663 |  |  | 904,876 |  |  | 199,872 |
|  | State I | State II | State III | State I | State II | State III | - |
| Final particle images (no.) | 56,631 | 44,856 | 31,119 | 109,001 | 112,932 | 56,528 | 108,876 |
| Map resolution (Å) | 3.8 | 3.7 | 3.9 | 4.2 | 4.3 | 4.5 | 3.5 |
| EMD accession code | 21824 | 21825 | 21826 | 21827 | 21828 | 21829 | 21830 |

## Supplementary Figures and Legends

**Supplementary Figure 1:**
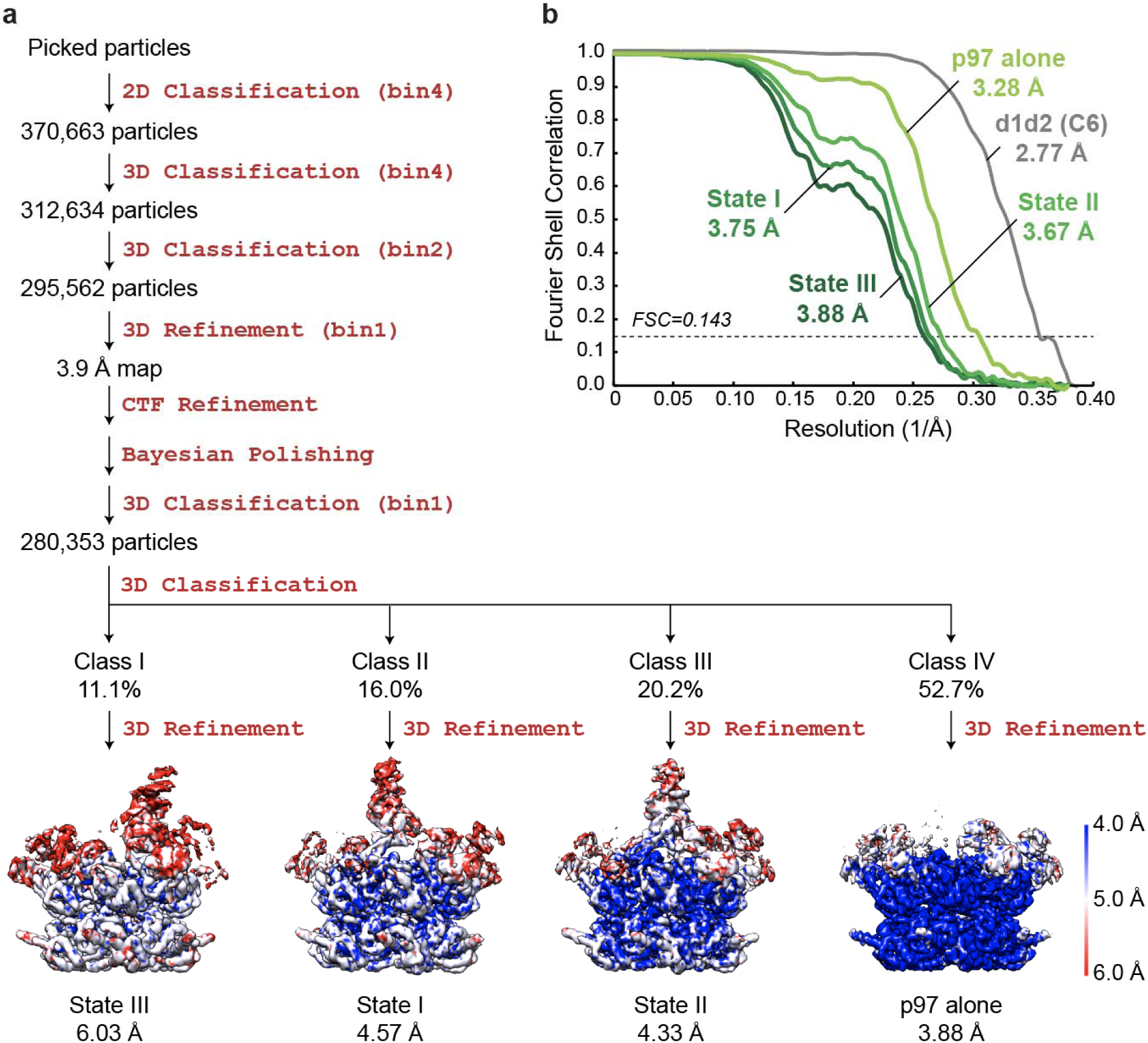
Single-particle cryo-EM analyses for the complex of p97 and Npl4/Ufd1. **a**, The workflow of data processing. The dataset was subjected to particle selection, 2D classification, and multiple rounds of 3D classification. Four maps including three different states of the complex and p97 alone were resolved. The maps were shown in similar orientation aligned in UCSF Chimera^39^. The local resolutions of the maps were calculated using LocalRes^37^. Resolutions after Relion 3D refinement and before post processing were listed. **b**, Fourier shell correlation curves of the masked maps after Relion post processing. The resolutions were determined by the FSC=0.143 criterion.

**Supplementary Figure 2:**
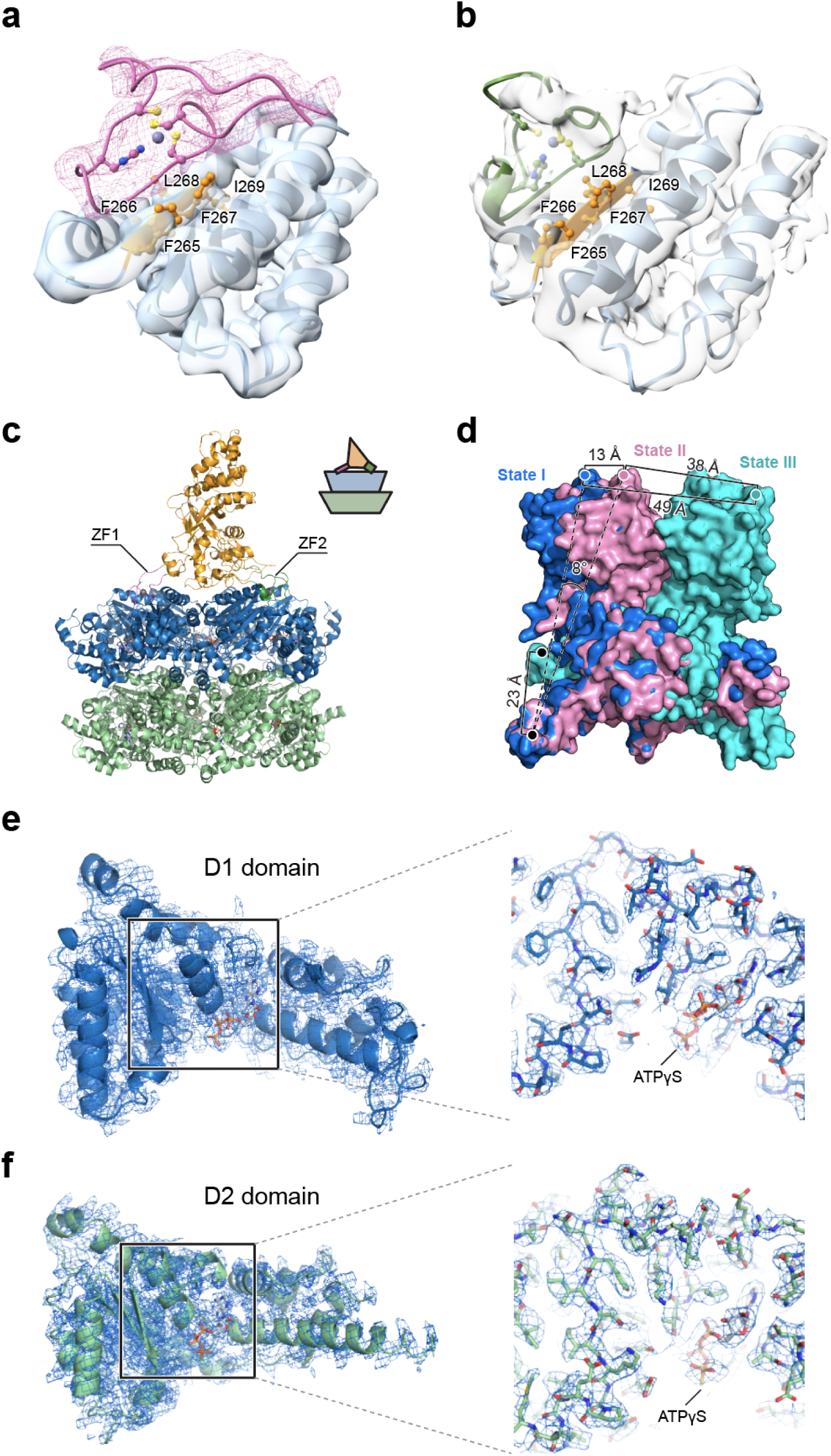
Interactions between the zinc finger motifs of Npl4 and p97. **a**, Interactions between ZF1 and the D1 domain of p97 (from State II of the p97-Npl4/Ufd1 complex). ZF1 forms a β-sheet with the large subunit of the D1 domain. The contacting β-strand in the D1 domain is colored in orange with the five hydrophobic residues (FFFLI) labeled. The map quality is good enough to enable model building of ZF1. **b**, Interactions between ZF2 and the D1 domain of p97 (from State III of the p97-Npl4/Ufd1 complex). ZF2 also forms a β-sheet with the large subunit of the D1 domain at the same location. The map quality is not sufficient to allow model building of ZF2, therefore a homology model was docked into the density as a rigid body. **c**, Structure of Cdc48-Npl4 complex from thermophilic fungi (PDB code: 6CHS). The position of Npl4 relative to Cdc48 is illustrated in the cartoon. Note that both zinc finger motifs are interacting with the D1 ring of Cdc48. **d**, Measurements of conformational changes of Npl4 from the three states. The structures were aligned based on the D1 and D2 rings of p97. **e**, A zoom view of the D1 domain showing the map around ATPγS. **f**, A zoom view of the D2 domain showing the map around ATPγS. Same color code is used as in Fig. 1.

**Supplementary Figure 3:**
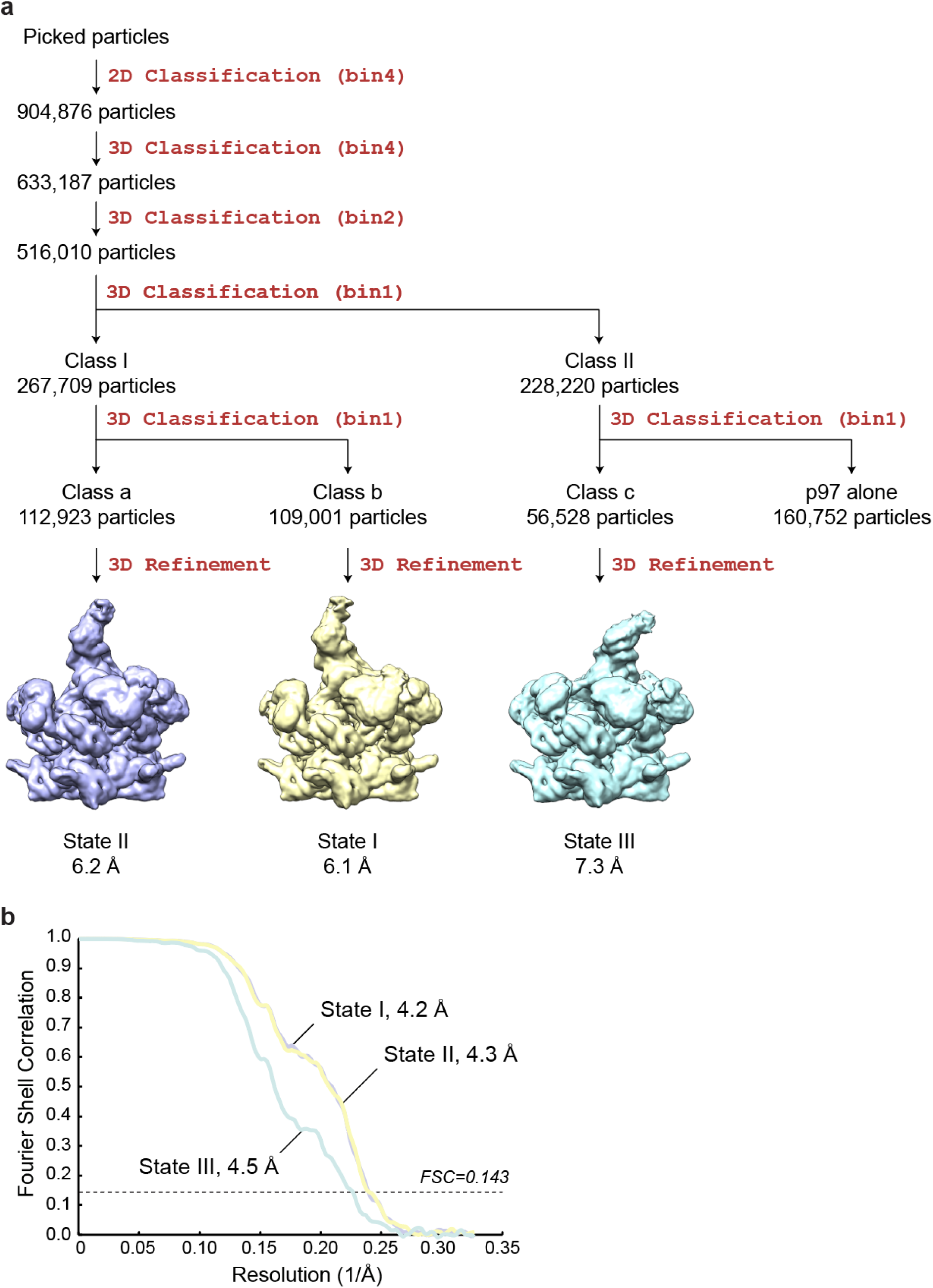
Single-particle cryo-EM analyses for the complex of p97, Npl4/Ufd1, and ubiquitinated Ub-Eos. **a**, The workflow of data processing. The dataset was subjected to particle selection, 2D classification, and multiple rounds of 3D classification. Three different states of the complex similar to the complex of p97 and Npl4/Ufd1 were resolved. The maps before Relion Post Processing were shown in similar orientation aligned in UCSF Chimera^39^. Resolutions after Relion 3D refinement and before post processing were listed. **b**, Fourier shell correlation curves of the masked maps after Relion post processing. The resolutions were determined by the FSC=0.143 criterion.

**Supplementary Figure 4:**
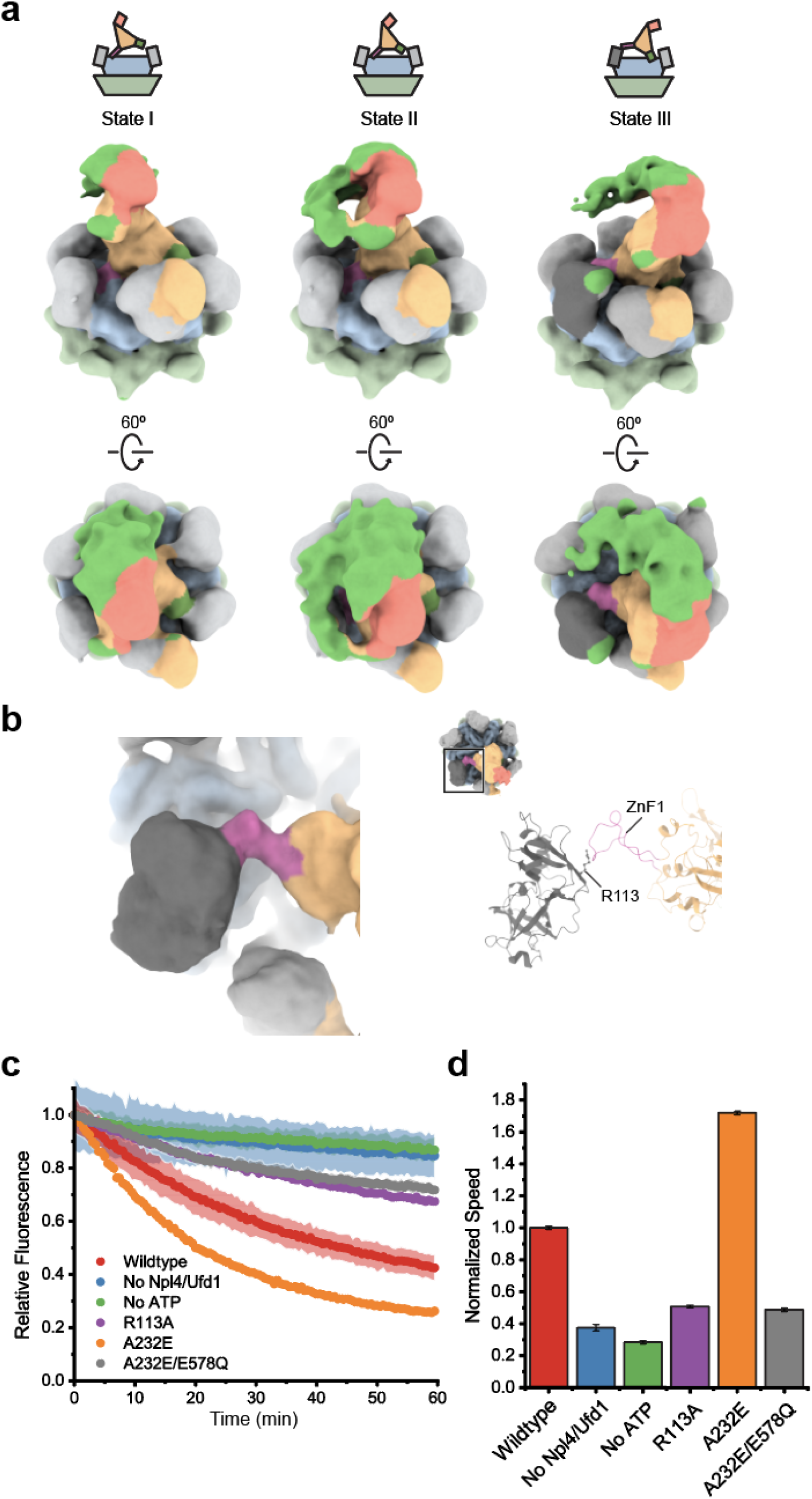
Single-particle cryo-EM structure of the complex of p97, Npl4/Ufd1, and ubiquitinated UbEos. **a**, Low pass filtered (15 Å) maps correspond to those in Fig. 2 shown at a lower threshold. Extra density (bright green) represents the polyubiquitinated Ub-Eos but was not well resolved due to the flexibility. **b**, The interactions between ZF1 and the N domain of p97 (from State III of the p97-Npl4/Ufd1-UbEos complex). The resolution is not sufficient to allow the fitting of side chains, therefore the crystal structure of the N domain and the ZF1 derived from panel a was docked into the density as rigid bodies. A tentative interacting residue R113 from the N domain was later mutated to test the relevance of the interaction. **c**, The unfolding activity of wild-type p97 and the mutants. **d**, The initial velocity corresponds to panel **e**. The error bands and bars in panels **c** and **d** represent the standard deviation from triplicate experiments.

**Supplementary Figure 5:**
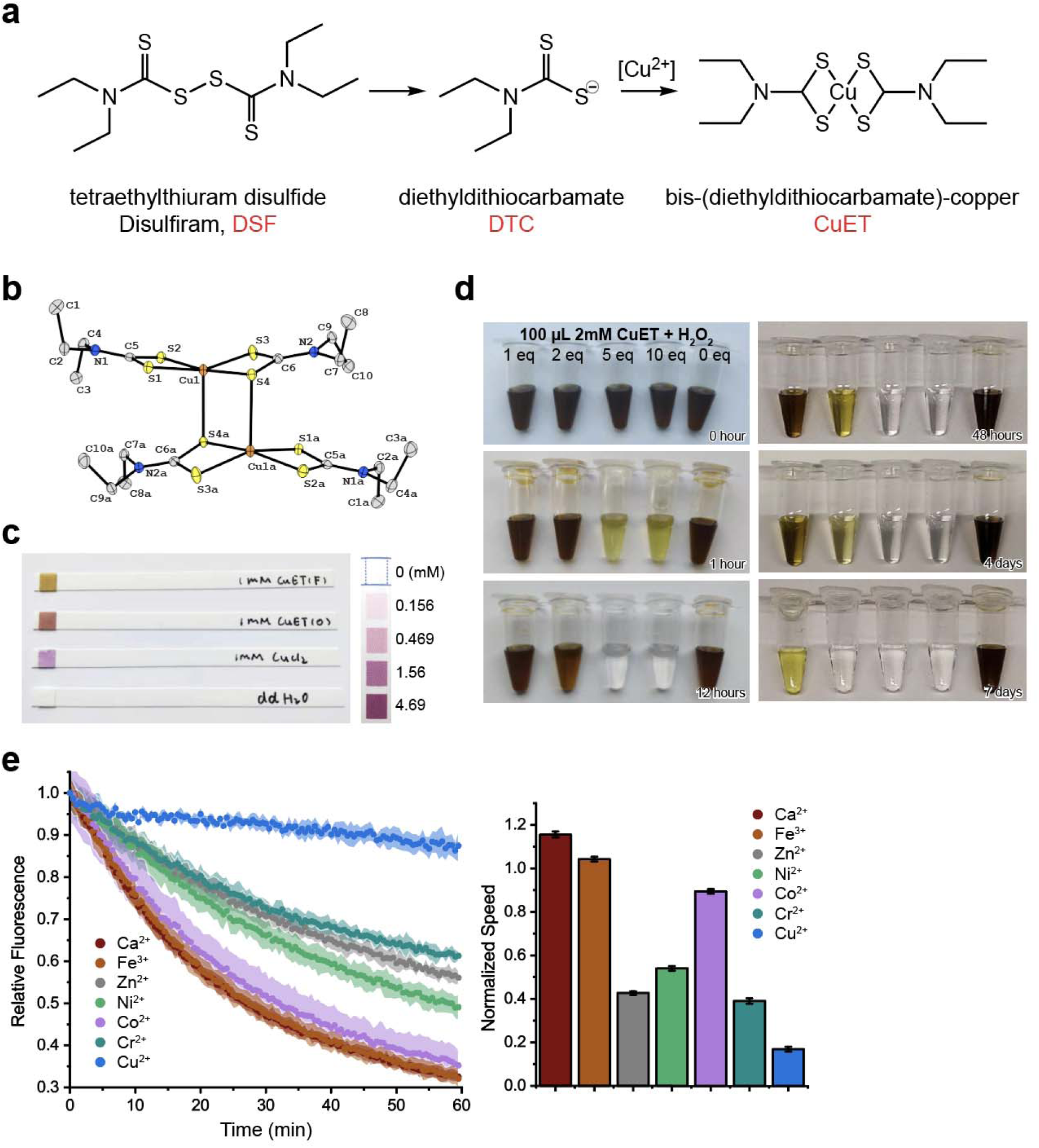
Disulfiram derivative CuET releases cupric ion in oxidative condition. **a**, Anti-alcohol abuse drug disulfiriram (tetraethylthiuram disulfide, DSF) is metabolized to bis-(diethyldithiocarbamate)-copper (CuET) in the liver^8^. **b**, The crystal structure of CuET synthesized for this study. **c**, Fresh CuET (F) solution in DMSO is brown colored. Old CuET (O) solution (> 7 days at 4 °C) releases cupric ions as shown by the copper strip with purple tint (Quantofix, Germany). **d**, CuET solution turns clear with additional hydrogen peroxide that accelerates the copper release in a concentration and time dependent manner. **e**, The unfolding activity and initial velocity of p97 in the presence of various metal ions at 25 μM. The error bands and bars represent the standard deviation from triplicate experiments.

**Supplementary Figure 6:**
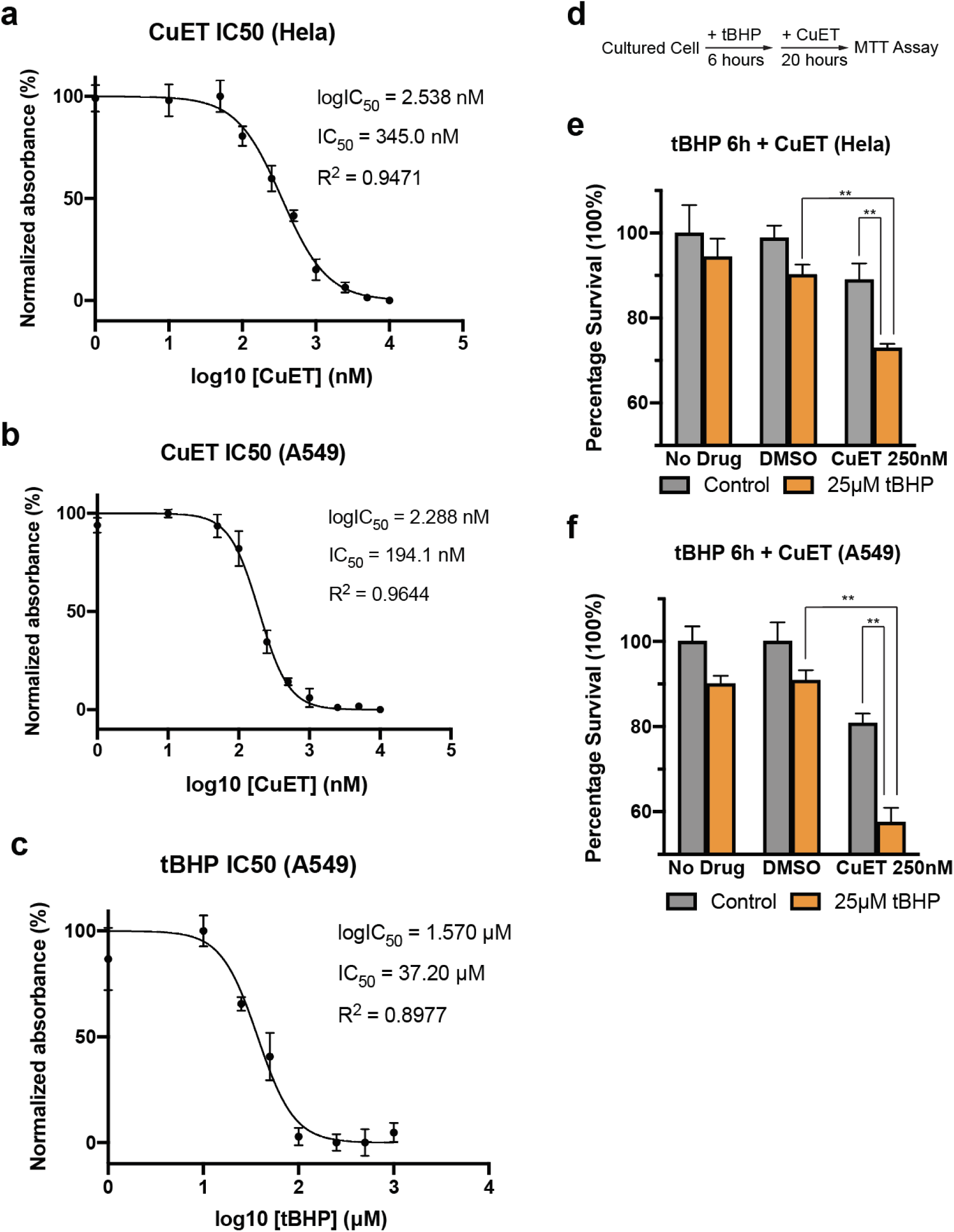
The toxicity of CuET is enhanced by oxidative stress in cultured cancer cells. **a, b**, The dose-response curves of Hela (**a**) and A549 (**b**) cells treated with CuET. Cell viability under various doses of CuET was measured by MTT assay after 48 hours. The CuET concentration at which cell viability reaches 50% (IC50 values) was determined by non-linear fit in Prism (GraphPad). **c**, The dose-response curve of A549 cells treated with oxidizing agent tert-Butyl hydroperoxide (tBHP). Cell viability was measured after 22 hours. **d-f**, Pre-treatment of tBHP to introduce oxidative state in the cultured cancer cells significantly enhanced the toxicity of CuET. **d**, Experimental scheme of evaluating CuET in oxidative conditions. **e, f**, Hela (**e**) and A549 (**f**) cells were treated with 25 μM tBHP for 6 hours before treated with 250nM CuET for 18 hours. Cell viability was measured by MTT assay. (n = 4 biological replicates, mean ± SEM, ** p<0.01, student T-test)

**Supplementary Figure 7:**
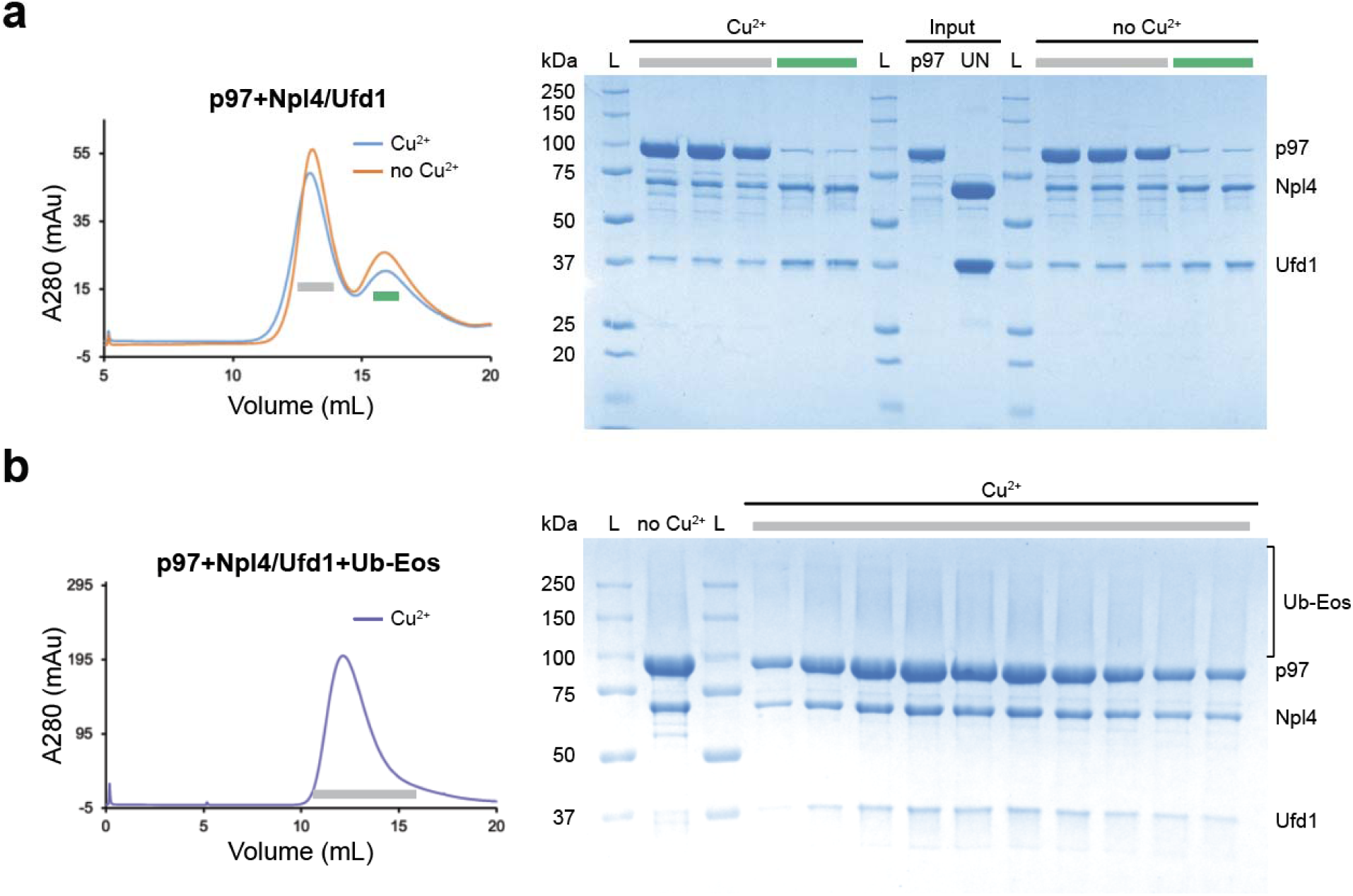
Cupric ions do not affect the complex formation of p97, Npl4/Ufd1, and polyubiquitinated Ub-Eos. **a**, The size exclusion chromatography profiles and SDS-PAGE gel after incubating p97, Npl4/Ufd1 in the presence (molar ratio 1:3:30) or absence of cupric chloride on ice for one hour. **b**, The size exclusion chromatography profile and SDS-PAGE gel after incubating preformed p97-Npl4/Ufd1 complex and polyubiquitinated UbEos in the presence of cupric chloride (molar ratio 1:2:10) on ice for one hour.

**Supplementary Figure 8:**
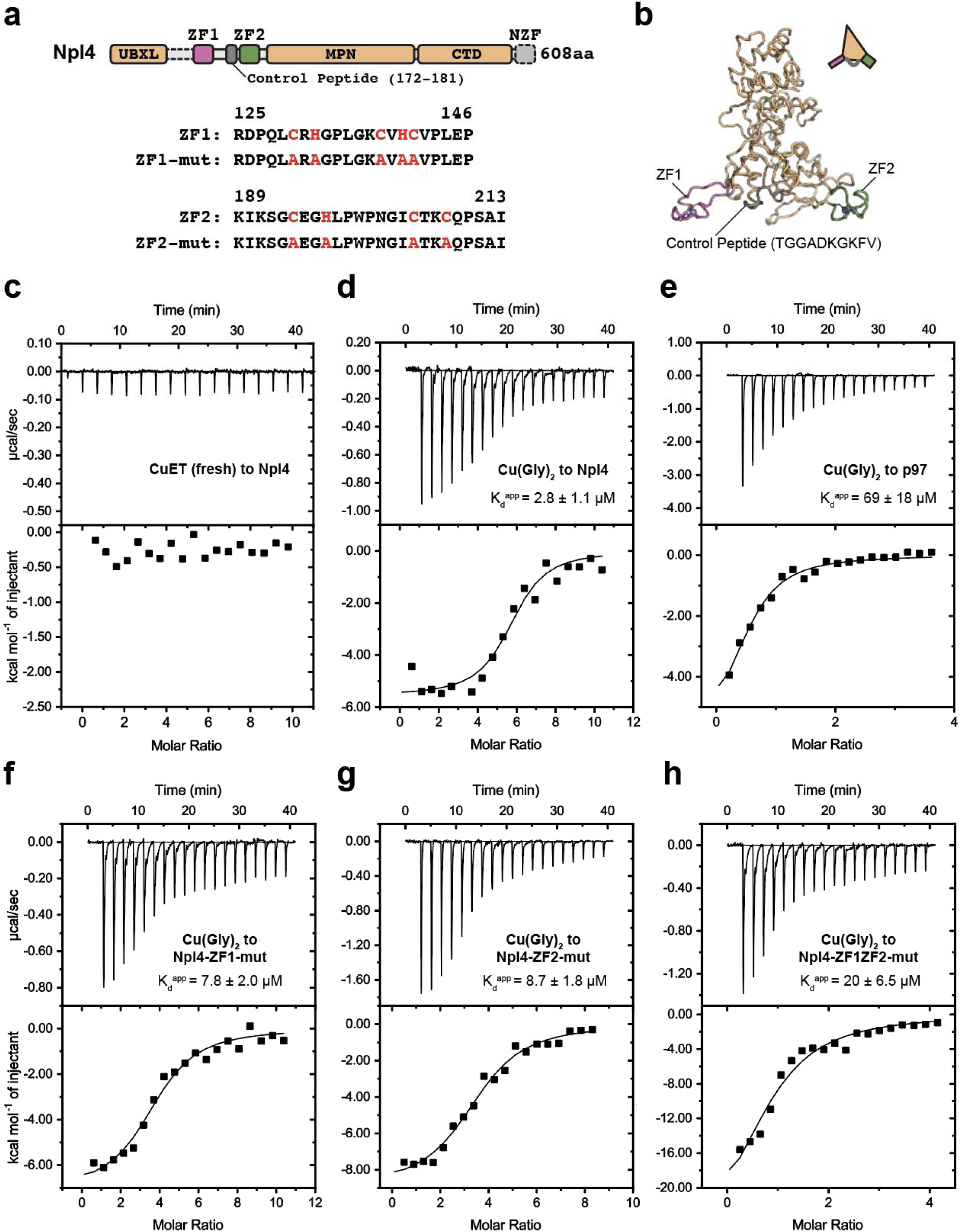
Cupric ions interact with the zinc finger motifs of Npl4. **a**, The zinc finger motif mutants used in this study. **b**, A homology model of human Npl4 derived from thermophilic fungi homologue (PDB code: 6CDD) using SWISS-MODEL^38^. Three peptides, ZF1 (magenta), ZF2 (green), and control peptide (grey, residues 172-181) were synthesized for the unfolding assay in Fig. 4. **c-h**, Isothermal titration calorimetry experiments. **c:** 1 mM CuET (fresh) titrated to 20 μM Npl4; **d:** 1 mM Cu(Gly)_2_ titrated to 20 μM Npl4; **e:** 5 mM Cu(Gly)_2_ titrated to 29 μM p97; **f:** 1 mM Cu(Gly)_2_ titrated to 20 μM Npl4-ZF1-mut; **g:** 1mM Cu(Gly)_2_ titrated to 25 μM Npl4-ZF2-mut; **h:** 0.8 mM Cu(Gly)_2_ titrated to 40 μM Npl4-ZF1ZF2-mut. Note that only apparent dissociation constant Kd^app^ are given as chemical reactions were taking place during the titration.

**Supplementary Figure 9:**
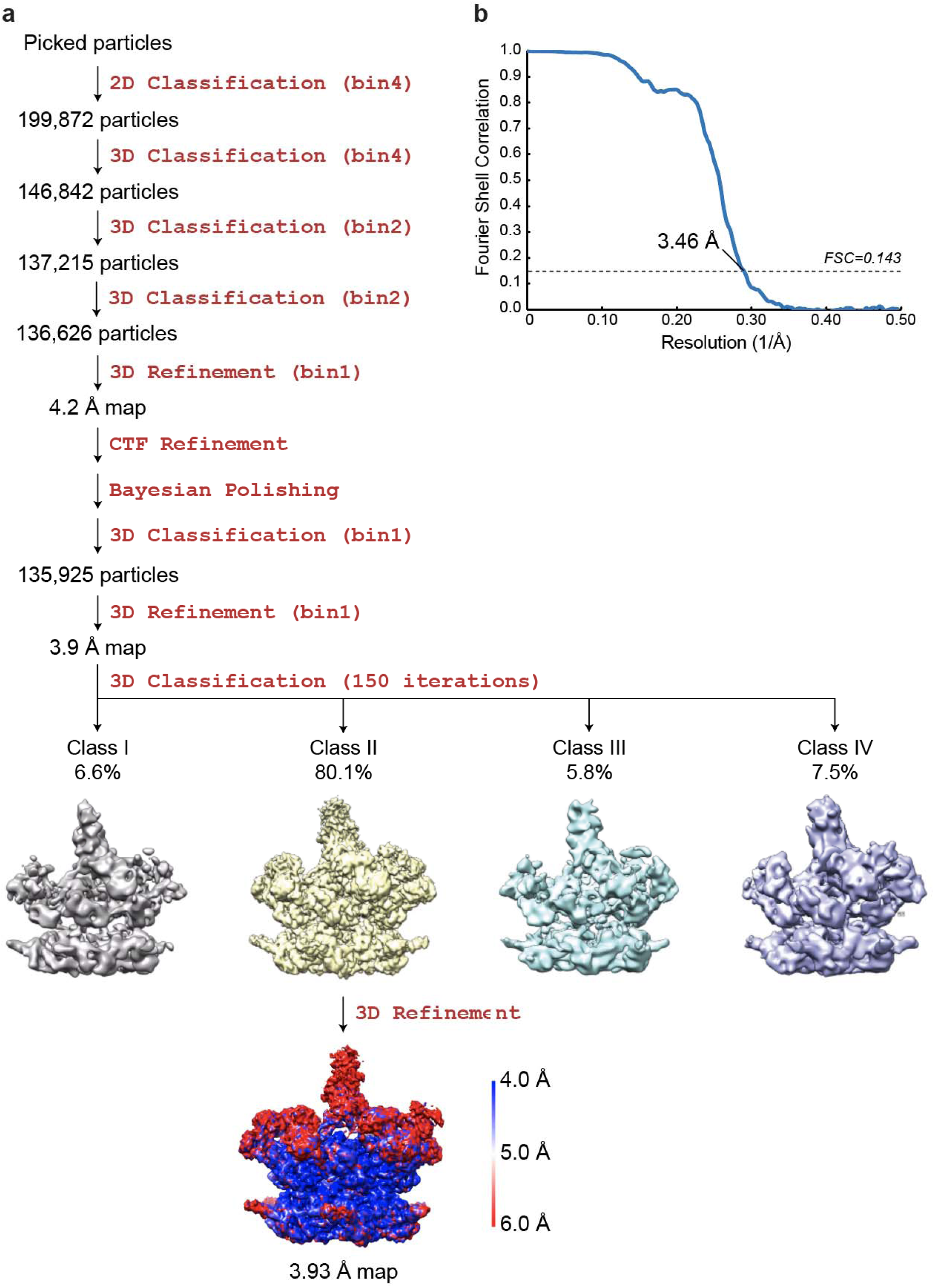
Single-particle cryo-EM analyses for the complex of p97 and Npl4/Ufd1 in the presence of cupric ion. **a**, The workflow of data processing. The dataset was subjected to particle selection, 2D classification, and multiple rounds of 3D classification. The maps after the last round of 3D classification were shown. All four maps are close to State II of the complex in the absence of cupric ion. The local resolution of the refined map was calculated using LocalRes^37^. The resolution after Relion 3D refinement and before post processing was listed. **b**, Fourier shell correlation curves of the masked map after Relion post processing. The resolution was determined by the FSC=0.143 criterion.

## Notes

### Competing Interest Statement

The authors have declared no competing interest.

### Summary of Updates

We have updated an error in figure Legend 5

